# Are members of the *Anopheles fluviatilis* complex conspecific?

**DOI:** 10.1101/2021.08.04.454551

**Authors:** Om P Singh, Ankita Sindhania, Gunjan Sharma, Shobhna Mishra, Surya K Sharma, Piyoosh K Singh, Manoj K. Das

## Abstract

*Anopheles fluviatilis sensu lato*, a primary malaria vector in India, was identified to be comprised of four cryptic species, provisionally designated as species S, T, U and V. However, Kumar *et al*. (Mol Ecol Resour, 2013;13:354-61) considered all of the then known three members of this species complex (S, T and U) conspecific. The specific status of species S and T was refuted based on the lack of sufficient barcode gap in mitochondrial-*CO1* and the perceived presence of heterozygotes in populations as detected through one of the two species-specific PCR assays employed for the cryptic species identification. The existence of species U was refuted claiming that earlier investigations have already refuted their existence. This conclusion is concerning because of the differential public health implications of members of the Fluviatilis Complex. Here we discuss problems associated with the *CO1*-based barcode approach for delimitation of cryptic species, the perceived heterozygosity between species S and T based on a species-specific PCR assay, and interpretation of published reports. We demonstrated that fixed differences do exist in the ITS2-rDNA sequence of species S and T with no evidence of heterozygotes in sympatric populations and, that the observed heterozygosity by Kumar *et al*. in the ITS2-based species diagnostic PCR is due to the high mispriming tendency of the T-specific primer with species S. We infer that mitochondrial DNA-based ‘barcoding gap’, an arbitrary threshold recommended for species delimitation, alone, is inadequate to delimit the members of *An. fluviatilis* complex.

---

*Anopheles fluviatilis s*.*l*. is one of the major malaria vectors in India which was recognized as a complex of four cryptic species, species S, T, U and species V (Subbarao *et al*., 1994; Nanda *et al*., 2013). The recognition of cryptic species in this species complex bears epidemiological significance owing to the contrast difference in host-feeding preference—an important determinant of malaria vectorial competence. Species S is highly anthropophagic (>91 percent) while species T, U and V are zoophagic (>87%) (Nanda *et al*., 1996; 2012). Species S was found to be an efficient malaria vector in forested areas in several states (Nanda *et al*., 2000, 2012; Sharma *et al*., 2006; Tripathy *et al*., 2010, Sahu *et al*., 2011), while species T and U were not found with sporozoite positivity (Sharma *et al*., 1995, Shukla *et al*., 1998), except for a few positive specimens of species T in Madhya Pradesh (Singh *et al*., 2015). Detailed bionomics of species V has not been studied yet. Species-specific differences were found in the nuclear second internal transcribed spacer (ITS2) as well as in the 28S ribosomal DNA (rDNA) (Manonmani *et al*., 2001; Singh *et al*., 2004, 2020; Chen *et al*., 2006), which were exploited for the development of PCR-based assays for their identification (Manonmani *et al*., 2001; Singh *et al*., 2004, 2020). However, the existence of species S, T, and U in the Fluviatilis Complex has been refuted by Kumar *et al*. (2013). Species S and T were considered conspecific based on the lack of the sufficient ‘barcoding gap’, an arbitrary threshold to delimit species, in the partial *cytochrome c oxidase subunit 1* locus (*CO1*) and observed presence of heterozygotes as detected through one of the two rDNA-based species-specific PCR employed for the species identification. The specific status of the third species, species U, was refuted based on misrepresentation of the published reports. Such a report has created confusion about the specific status of cryptic species and hampered further research on the differential bionomics of members of the species complex.

DNA barcoding has emerged as a global standard for species identification which, in the case of animals, utilizes the sequence divergences at ∼600 bp mitochondrial gene *cytochrome oxidase 1* (*CO1*) for the discrimination of the closely allied species. The advent of the barcoding system has discouraged to some extent conventional methods to delimit species based on the evidences pointing to genetic isolation in cryptic species. While barcode has been successfully used for the delimitation of species in a wide array of animal taxa, it failed to delimit the species in many cases where a sufficient gap does not exist due to the incomplete lineage sorting and introgression of mitochondrial DNA. Although the use of multiple markers (nuclear and mitochondrial) has been advocated to overcome this problem owing to frequent reports of mito-nuclear discordance, but such an approach is less frequently used. In the case of *An. fluviatilis*, Kumar *et al*. barcoded species S and T and considered them conspecific merely on the basis of lack of sufficient barcoding gap between species S and T without considering fix differences in nuclear rDNA (Manonmani *et al*., 2001; Chen *et al*., 2006; Singh *et al*., 2006) and differences in host-feeding preference (Nanda *et al*., 1996, 2012).

Kumar *et al*. considered differences in rDNA sequences “just ribosomal DNA genetic variants” which “may not have any taxonomic significance”. We argue that fixed and conserved differences in rDNA, especially in a sympatric population, are clear evidence of genetic isolation. However, due to limited data prevailing in public domain on ITS2, we generated ITS2-rDNA sequences of *An. fluviatilis* specimens collected from a forested belt under Odisha and West Singhbhum hills (see **Supplementary file S1**) where species S and T have been reported to be sympatric. The analysis of ITS2 sequences (species S=106, T=46) confirmed two distinct and conserved types of sequences in morphologically identified *An. fluviatilis*, one identical to Species S (Singh *et al*., 2006) and the other identical to species T (Chen *et al*., 2006). The sequence alignment has been shown in **Figure 1**. The species S and T differ at a total of 14 nucleotide base positions (including 2 indels) with a genetic distance of 3.28% (Kimura 2-parameter). No intraspecific variation in DNA sequences was observed. In species T, there were mixed bases at nucleotide positions 91 and 198 (**Figure 1**) which were consistently present in all species T samples. We didn’t find any heterozygotes in these populations. In other words, these two sequences are fixed in a sympatric population. If we assume that the two molecular forms in respect to ITS2 are freely interbreeding without genetic constraints, we expect heterozygotes or evidence of homogenization of ITS2 sequence, that was absent. Two distinct and fixed ITS2 sequences in a sympatric population with a lack of homogenization or hybridization provide clear evidence of reproductive isolation between them.

**Figure 1.**
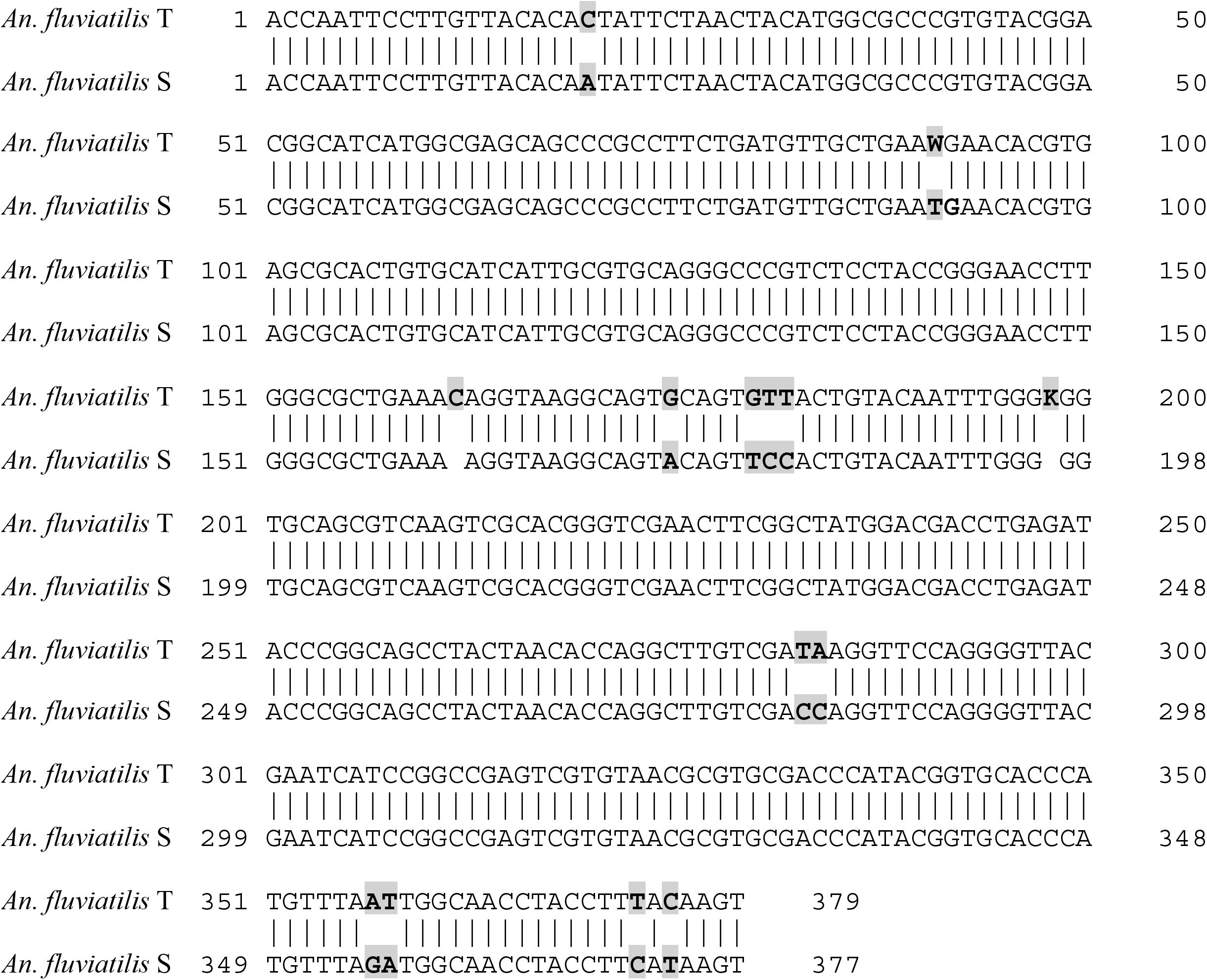
Sequence alignment of species ITS2-rDNA of species S and T. The differences in nucleotide sequences are highlighted.

Another argument by Kumar *et al*. in the favour of conspecificity of species S and T is the perceived presence of heterozygotes of species S and T in the population. Kumar *et al*. observed the presence of faint species T-diagnostic bands along with prominent S-specific bands in an ITS2-based species diagnostic PCR assay (ITS2-ASPCR) (Manonmani *et al*., 2001) in samples that were identified as Species S by another diagnostic PCR based on 28S-rDNA (Singh *et al*., 2004) and considered them to be heterozygotes. However, Kumar *et al*. didn’t verify the presence of heterozygosity through DNA sequencing. We also observed the presence of faint T-diagnostic band (∼450 bp) in samples belonging to species S, in addition to the prominent S-diagnostic band (∼345 bp) (**Figure 2**). Faint T-specific band can be better seen when relatively a higher amount (10-15 µL) of PCR product is loaded in the well of agarose gel. However, when we sequenced ITS2 region of such samples, an unambiguous sequence corresponding to species S, without any trace of the T-specific sequence, was obtained (**Figure 3**). Had it been heterozygote, we expect mixed bases at all the nucleotide substitution positions and collapse of DNA sequence chromatogram beyond the point of indels, which were absent in all such samples. Thus, the presence of T-specific diagnostic band in species S can be either due to mispriming of T-specific primer (T-R) with species S or due to extremely low copy number of T-specific ITS2 due to the intragenomic sequence variations not detectable through DNA sequence chromatogram (Sharma et al., 2016). We, therefore, checked the priming efficiency of primer T-R on five clonal ITS2 products of species S (plasmid DNA with ITS2 insert) of known sequences which were confirmed to have species S-specific ITS2 sequence and were free from PCR error. PCR was performed with primers 5.8S-F (universal primer from 5.8S rDNA) and T-R used by Manonmani et al. (2001), in stringent PCR conditions (see **Supplementary file S1**). We observed a prominent band (**Figure 4**) corresponding to the diagnostic band size of species T in all clonal samples confirming the high mispriming tendency of the primer. To confirm that this band is not due to the contamination of reagents or carryover of DNA or aerosol DNA, we sequenced two PCR products amplified from clonal samples. We obtained a clean sequence that was identical to species S (except in the T-R primer region which is derived from the primer) (**Figure 5**). This exhibits that T-specific primer has the tendency of mispriming with species S template. In another experiment, we also checked mispriming of T-R on *An. minimus s*.*s*. samples, which have an identical sequence to species S in the primer (T-R) region. All the *An. minimus* samples (n=10) tested with ITS2-ASPCR, provided false T-diagnostic band ignoring mismatches in primer (**Figure 2**). We also designed and carried out an RFLP assay to check the presence of heterozygote in species S. Assuming that restriction enzymes have tremendous specificity to their recognition site, we designed species T-specific RFLP using enzyme *Bts*I*-v2*. The *An. fluviatilis* S samples including those with the observed heterozygosity in ITS2-ASPCR were subjected to this PCR-RFLP assay. All the samples (n=27) with observed heterozygosity in PCR assay didn’t cleaved with *Bts*I*-v2* (**Figure 6)**. These experiments show that the observed heterozygosity in ITS2-species diagnostic assay may be due to mispriming of species T-specific primer with species S template. We, therefore, ruled out the possibility of the presence of heterozygotes based on ITS2-ASPCR until this is not proved through DNA sequencing.

**Figure 2.**
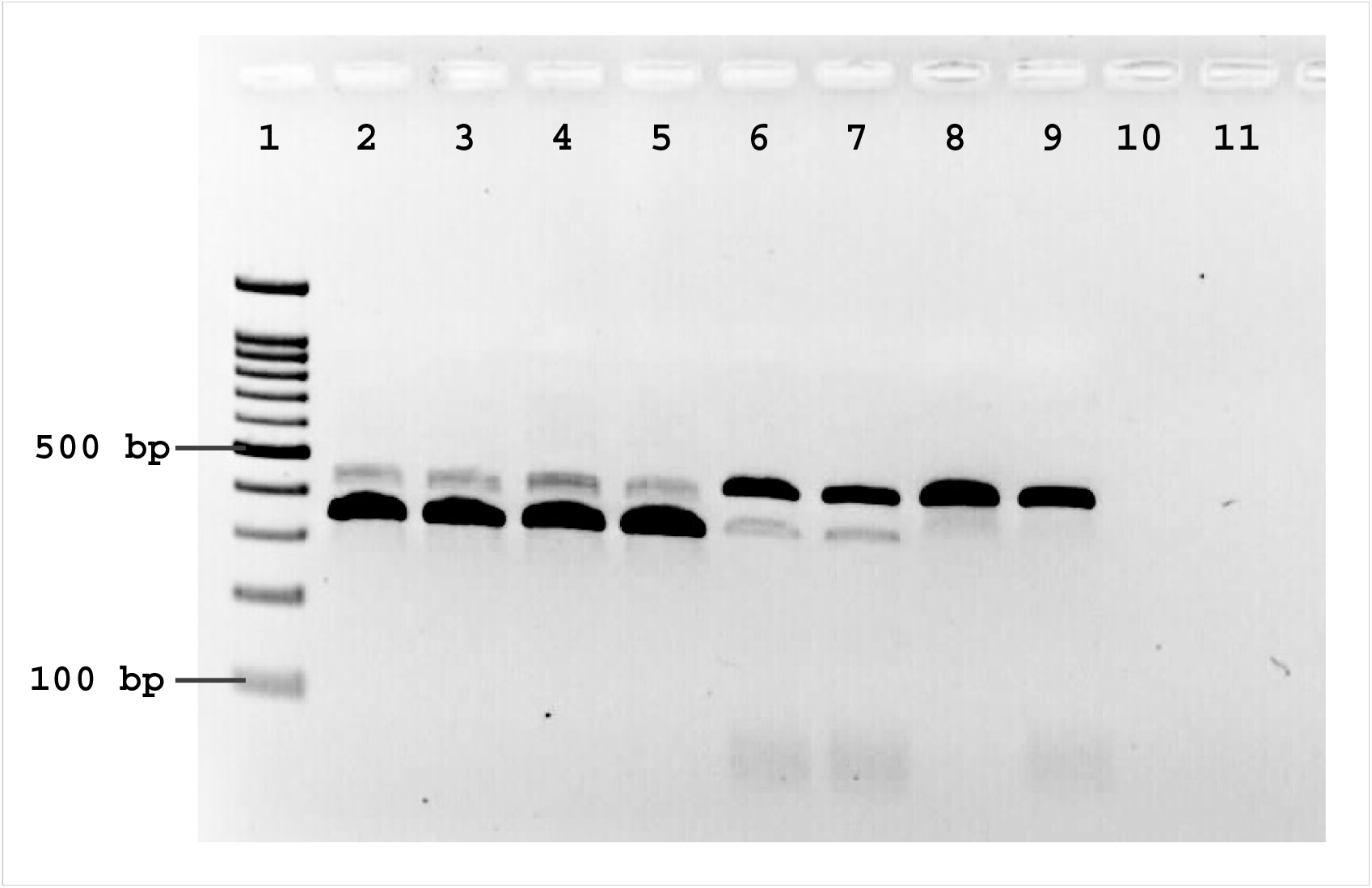
Species-diagnostic PCR (Manonmani *et al*., 2001) showing cross-reactivity of T-R. with species S and *An. minimus s*.*s*. The size of species T- and S-specific amplicons are 345 and 450 bp, respectively. Lanes 2 & 3: species S, lanes 4 & 5: clonal S samples; lanes 6 & 7: *An. minimus*, lanes 8 & 9: species T; lane 1: 100 bp DNA ladder, lane 10: negative control without DNA, lane 11: negative control with pGemT easy vector.

**Figure 3.**
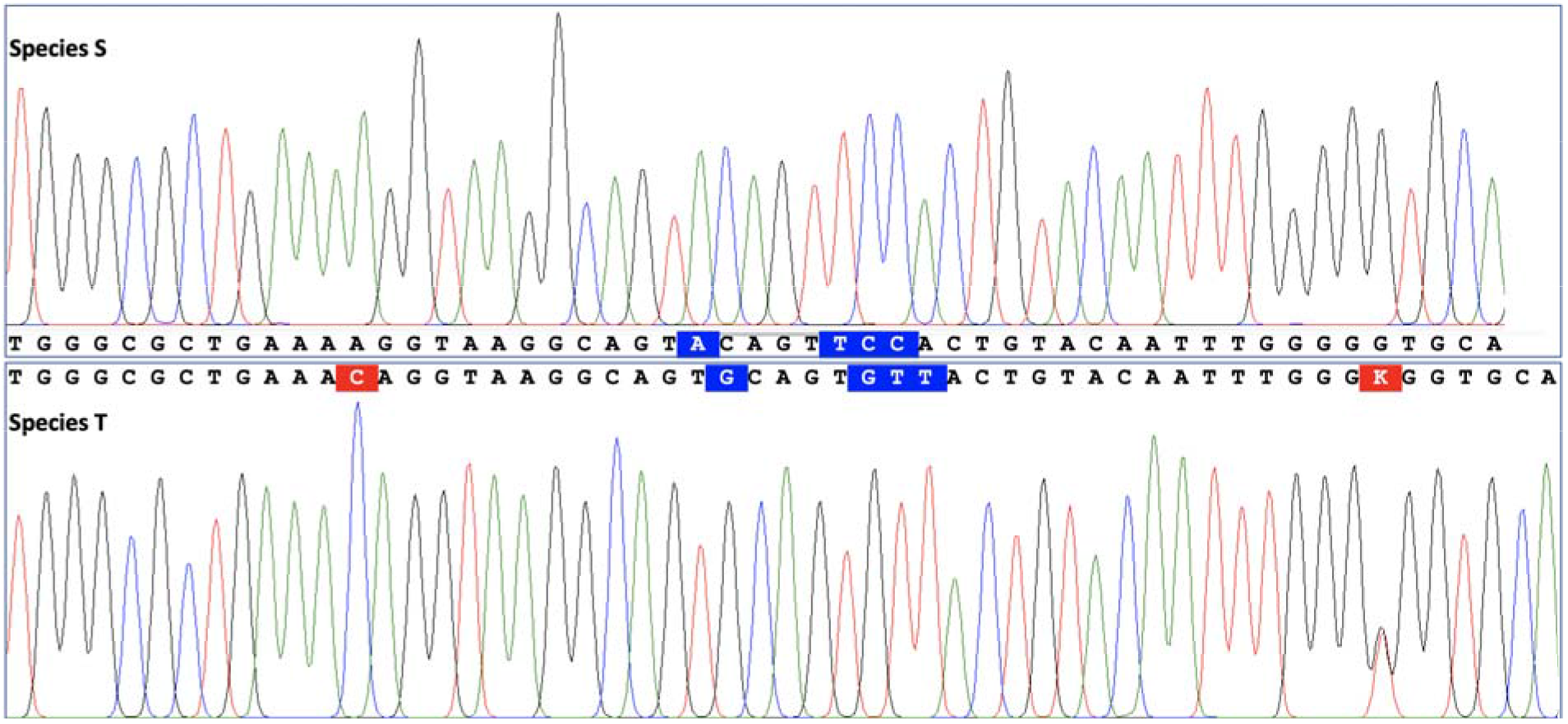
DNA sequence chromatograms of a portion of ITS2-rDNA of species S and T showing nucleotide sequence and length polymorphism. The species S sequence shown here belongs to a sample that showed the presence of both species S and T diagnostic amplicon in ITS2-based species diagnostic assay. It may be noted that there is no signature of heterozygote in the DNA sequence. The nucleotide differences in sequences are highlighted (indels with red colour and substitutions with blue colour).

**Figure 4.**
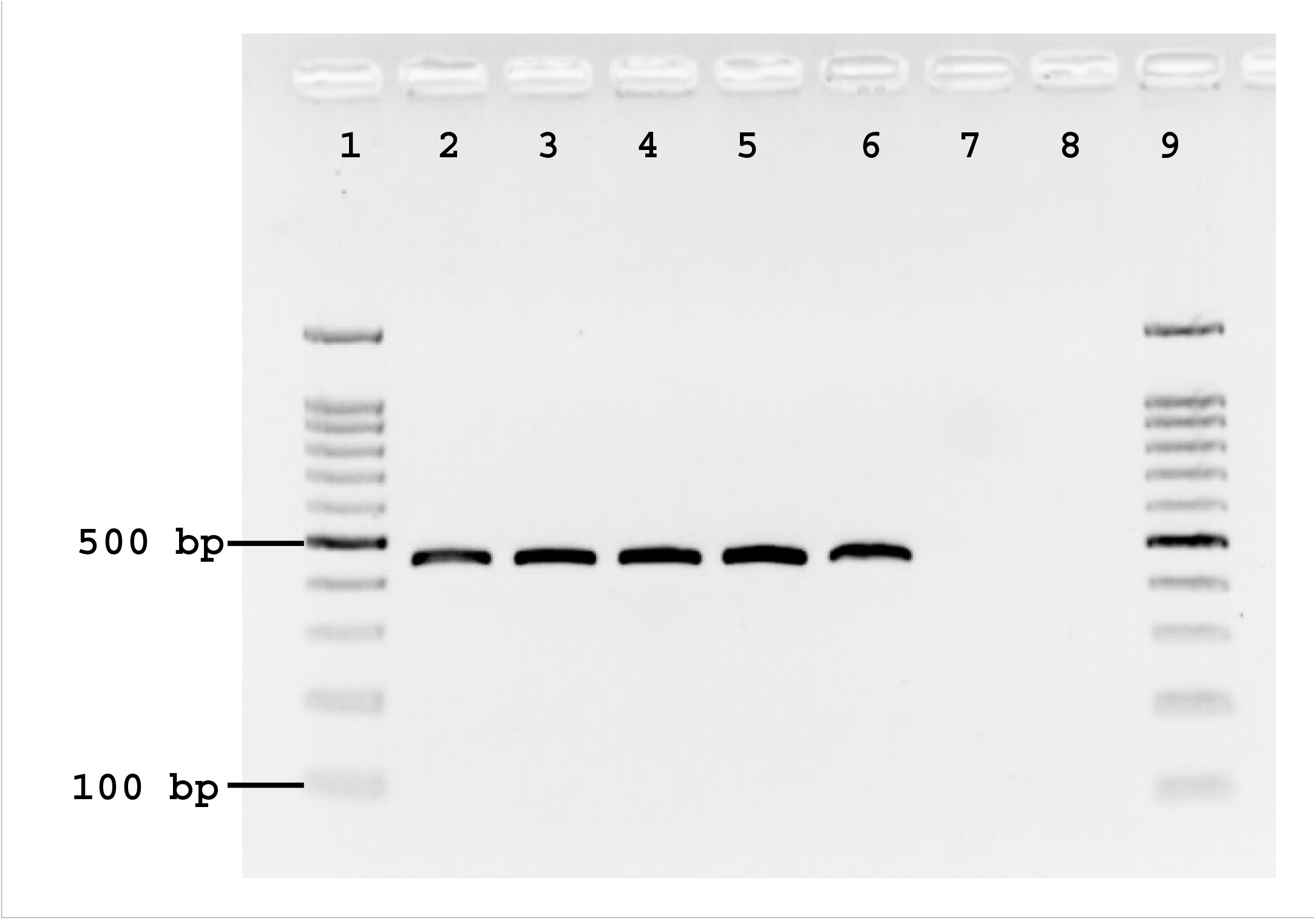
PCR products (450 bp) amplified with primers 5.8S-F and T-R (T-specific) on clonal samples of species S (ITS2) showing mispriming of T-R with species S template. Lanes 1 & 9: 100 bp DNA ladder; lanes 2—6: clonal sample of species S; lane 7: negative control with DNA, lane 8: negative control with pGemT easy vector.

**Figure 5.**
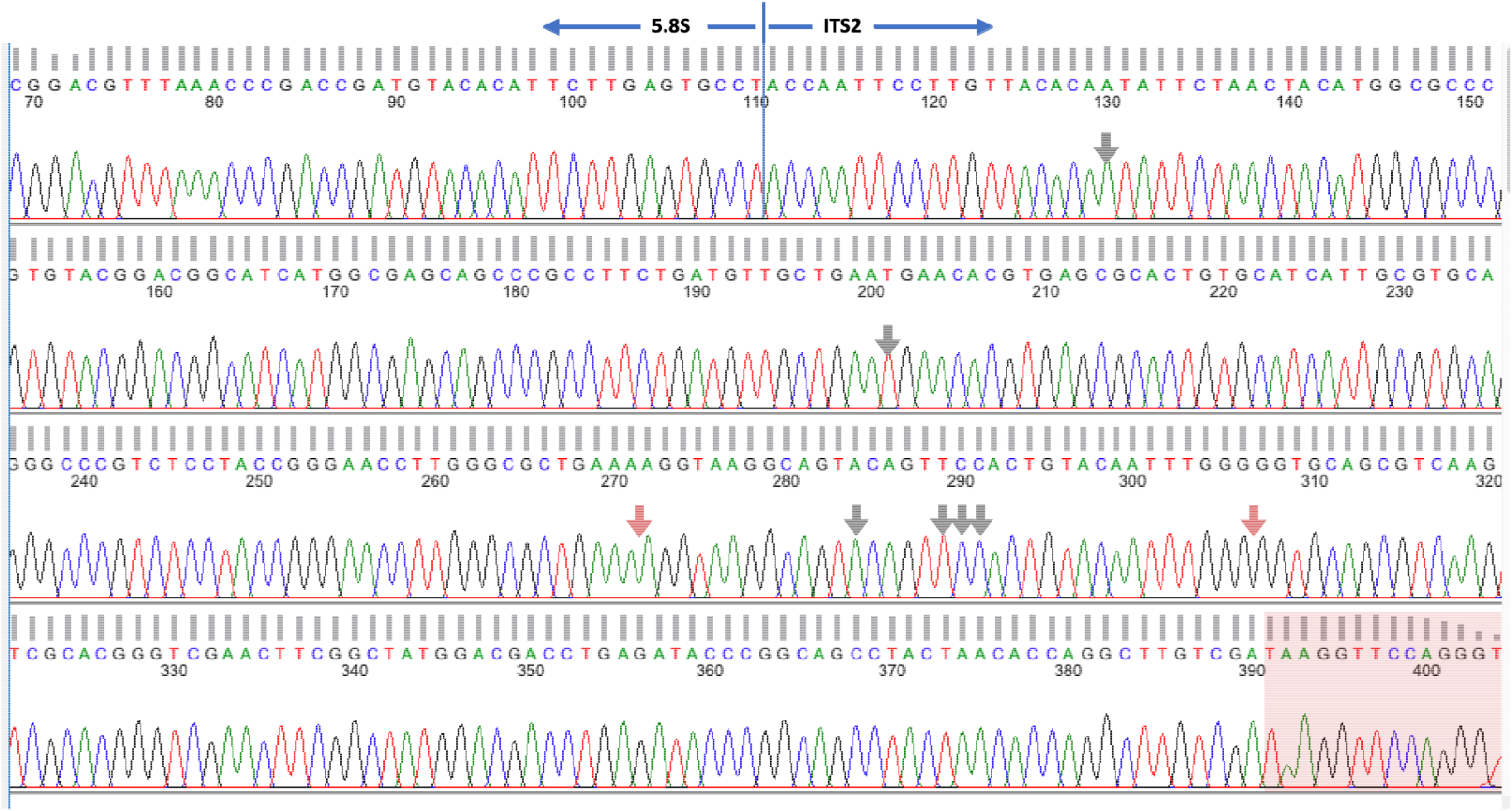
DNA sequence chromatogram of non-specific primer extension product amplified with universal primer 5.8SF and T-specific primer (T-R) on species S clonal template. The amplified product is identical to species S sequence except in primer region (highlighted with pink colour) which is derived from T-R primer. Vertical arrows indicate base positions where species S and differs from species T (grey arrow=substitution; orange arrow=indel)

**Figure 6.**
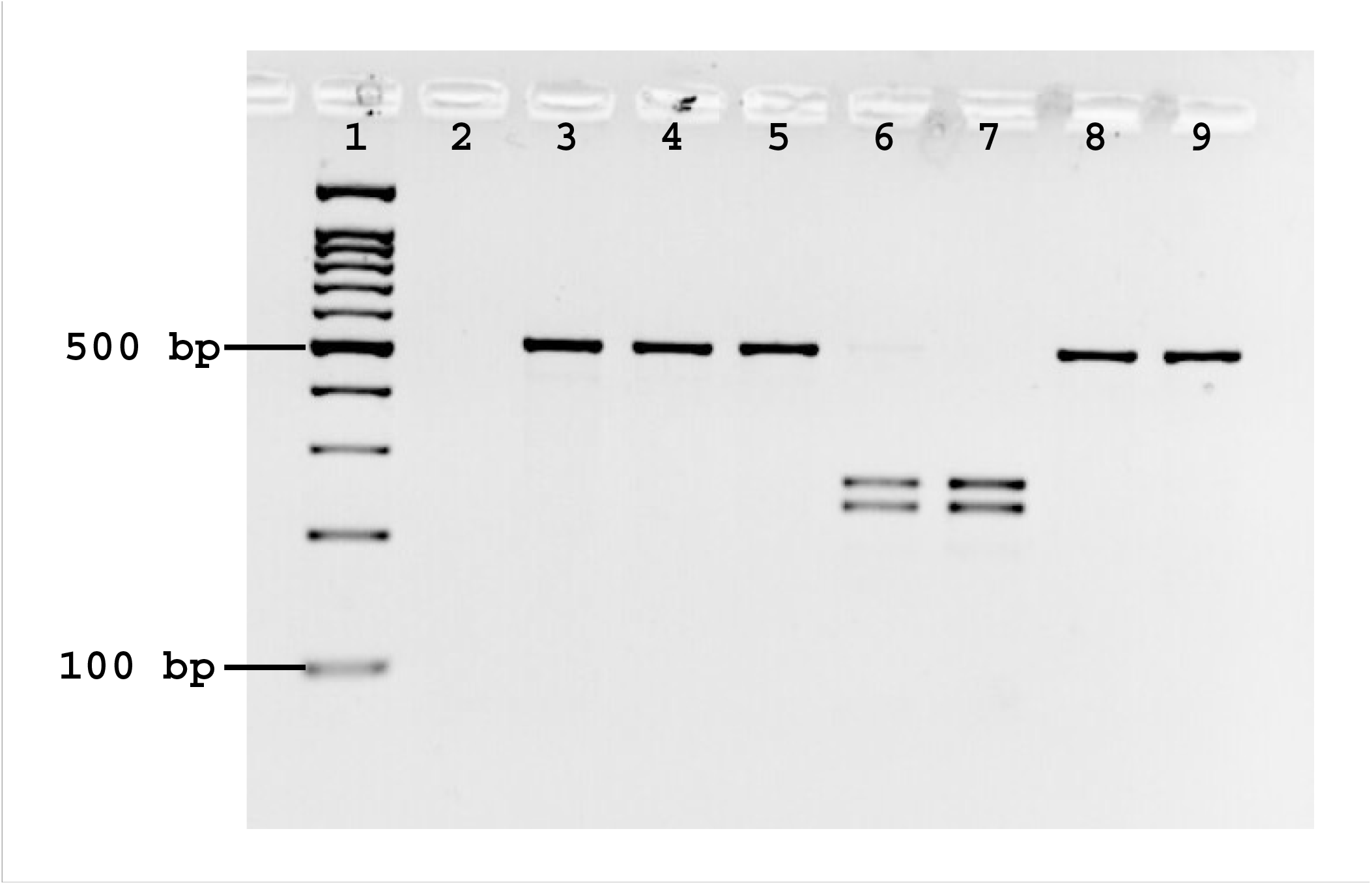
Results of PCR-RFLP (*Bts*I*-v2*) on species S and T samples. The PCR products amplified with ITS2A and ITS2C were subjected to restriction digestion. Lane 1: 100 bp ladder; lane 2: negative control, without DNA; lanes 3—5: species S; lanes 6 & 7: species T; lanes 8 & 9: species S and T, respectively. without enzyme treatment.

Another plausible and strong evidence supporting the distinct specific status of species S and T is the contrast differences in their host preferences. Species S has been found mainly anthropophilic (>91%) whereas species T was zoophagic (>87%) in sympatric populations (Nanda *et al*., 1996; 2012).

Kumar *et al*. also refuted the existence of species U without any supporting data generated by themselves. Albeit, the argument for such refutation is based on an erroneous interpretation of previous findings. Kumar *et al*. stated “*earlier investigations have refuted the existence of species U*” without citing any reference of the published report. However, in the latter part of their paper, they referred to Chen *et al*. (2006) in support of their statements. Citing Chen’s reference, the authors stated “… *however, genetic introgression between species U and species T had been described in some parts of India (Chen et al. 2006)”*. This is not in agreement with the statement by Chen *et al*. (2006) concerning the Indian population.

Conversely, Chen *et al*. (2006) have clearly stated “*We sequenced the D3 region* (of 28S rDNA) *from 12 individuals of species T and eight of species U from Uttar Pradesh in north-western India and the sequences were consistently distinct, albeit for a single substitution*”. Chen *et al*. (2006) further concluded “*Therefore*, An. fluviatilis *T and U may be regarded as well-defined but very closely related species*”. Chen *et al*. (2006) also recorded differences of six bp in ITS2 between species T and U. The absence of heterozygotes through chromosomal (Subbarao, 1994) and molecular markers (Chen *et al*., 2006) in a population where species T and U are sympatric (Uttarakhand, undivided Uttar Pradesh) evident the presence of reproduction barrier between these two species. Moreover, differences in palpal ornamentation of sympatric species T and U have also been noted (Sharma et al, 2020) Introgression in these species, if any, is a subject of further research.

Kumar *et al*. also endorsed an already resolved controversy on the conspecificity of species S with *An. harissoni* prevailed in literature in the past (Garros *et al*., 2005; Chen *et al*., 2006) ignoring subsequent work by Singh *et al*. (2006). We would like to mention that the inference that species S is conspecific with *An. harissoni* was based on the circumstantial evidence of similarity in a 336 bp conserved stretch of 28S rDNA.

However, Singh *et al*. (2006) showed that significant differences do exist between these two species in extended 28S rDNA, ITS2 as well as in *cytochrome oxidase II*, with a high degree of divergence. Surprisingly, this report was ignored by Kumar *et al*. creating confusion among the reader. The across-the-board application of mere *CO1* sequences for delimitation of closely related species or recently diverged species is a subject of controversy. The proponent of barcoding of life claims that interspecific variation in partial *CO1*-mtDNA exceeds intraspecific variation by an arbitrary threshold called ‘barcoding gap”. Although DNA barcode has successfully been used for delimitation of species where sufficient gap exists between intra- and interspecific *CO1* sequence divergence, such gap often doesn’t exist in closely related species or recently diverged species with incomplete lineage sorting. Considerable overlap in the range of intra- and interspecific *CO1* sequence divergence has been well documented (Meyer & Paulay, 2005; Wiemers & Fiedler, 2007). It was noted that a high proportion of well-differentiated species has similar or even identical *CO1* sequences (Wiemers & Fiedler, 2007; Conflitti *et al*., 2012). Among 449 dipteran species, Meier et al., (2006) find a low success rate (< 70%) based on tree-based and newly proposed species identification criteria of the barcode system. In the case of butterflies, Only 77% of species could be accurately identified using the barcode data (Eliass *et al*., 2007). In mosquitoes, Carter *et al*. (2019) noted that *CO1* can not differentiate *An. gambiae* and sister species *An. arabiensis*. Even the whole mitochondrial genome of *An. arabiensis, An. gambiae* and *An. coluzzii* failed to show clear species division (Hanemaaijer *et al*., 2018). Several studies have reported discordant patterns between mtDNA and nuclear markers, commonly referred to as ‘mito-nuclear discordant’ (Toews & Brelsford, 2012; Després, 2019). It has been proposed that this discordance may be due to incomplete lineage sorting or introgression (Toews & Brelsford, 2012); however, some reports shows that the discordance cannot be explained by incomplete lineage sorting or introgression (Ivanov *et al*., 2018; Hinojosa *et al*., 2019). It has been proposed to use multiple nuclear and mitochondrial genes for delimitation of species (Vences *et al*., 2005; Beebe, 2018) and adopting integrative taxonomy approaching genetics, morphology, ecology, behaviour and geography (several references cited by er *et al*., 2018). Whatever method is used for species delimitation, one should consider important genetic principles lying with speciation, i.e., cessation of gene flow or genetic isolation which is a primary pointer of ongoing speciation (Bock, 2004). Additional data on molecular markers demonstrating gene flow restriction between S and T of the Fluviatilis Complex, crossing experiments, etc, is suggested in view of mito-nuclear discordance.

In conclusion, there is no plausible evidence so far indicating a breakthrough in reproductive isolation between *An. fluviatilis* species S and T so far in a sympatric population. The absence of a genetic barcode gap between species S and T may be due to retention of ancestral polymorphisms or possible introgression, which is not enough to prove synonymy in a condition when fixed differences in nuclear ITS2 do exist between these two species with no signature of hybridization. The inferred heterozygosity between species S and T through an ITS2 based species diagnostic assay by Kumar *et al*. is ruled out due to the high mispriming tendency of T-specific primer. Contrast difference in host-selection attributes further supports their independent biological entity. Further study is required to confirm the genetic isolation among all members of the Fluviatilis Complex, preferably in a sympatric population.

## Authors’ Contribution

OPS designed and executed research, wrote the first draft of the manuscript, AS, GS and SM performed PCR, cloning and sequencing, SKS, PKS and MKD carried out fieldwork and sampling of mosquitoes. All authors have read and approved the final version of the manuscript.

## Acknowledgements

The work was supported by the Indian Council of Medical Research (ICMR) project grant (6/9-7(17)/2013-ECD-II). AS was supported by the Department of Science & Technology fellowship, GS and SM were supported by the Indian Council of Medical Research Fellowship. Authors are grateful to Dr SK Subbarao for the critical review of the manuscript and constructive suggestions.

