## Supplementary file S1 for "Are members of the *Anopheles fluviatilis* complex conspecific?"

**Supplementary information S1**

**Material and Methods**

**Mosquito collection and processing**

Wild female *An. fluviatilis* and *An. minimus* specimens were collected from three forested localities from Odisha state, i.e., Kedunjhar (21.6°N 85.6°E), Koraput (18.8°N 82.7°E), and Sundergarh (22.1°N 84.0°E) districts, and a bordering district West-Singhbhum (22.4° N, 85.4° E) of Jharkhand state. Mosquitoes were morphologically identified, dried and transported to the laboratory at Delhi in individual microfuge tubes containing a piece of silica gel. DNA was isolated from morphologically identified *An. fluviatilis* and *An. minimus* using the method described by Livak (1984).

**DNA sequencing**

*An. fluviatilis* samples were PCR amplified using primers ITS2A (Beebe & Saul, 1995) and ITS2C (Singh *et al.,* 2010) designed from flanking 5.8S-rDNA and 28S-rDNA region, respectively. PCR was carried out using Hot Start Taq 2X Master Mix (New England Biolabs Inc) in a 20 µL reaction mixture containing 1.5 mM of MgCl_2_, 0.5 unit of Taq polymerase and 0.25 µM of each primer. The PCR cycling conditions included a denaturation step at 95 °C for 3 min, 35 cycles each with denaturation at 95 °C for 30 sec, annealing at 55 °C for 30 sec and extension step at 72 °C for 45 sec and final extension at 72 °C for 5 min. PCR products were cleaned using Exo-Sap and sequenced using ABI BigDye terminator v3.2 following the vendor’s standard protocol. Primers used for sequencing were ITS2A and ITS2C. A few *An. minimus* collected from Kedunjhar was found in sympatric association with *An. fluviatilis* in Kedunjhar were also sequenced for ITS2.

**Cloning and sequencing**

The specificity of T-R primer (Manonmani *et al.*, 2001) with gDNA from species S was tested on the pure clone of rDNA with known sequence harvested from species S. The purpose of choosing a clone as a PCR-template was to eliminate the possibility of PCR extension with any intragenomic variant copy of rDNA, if any present in gDNA, which may have a complementary sequence to T-specific primer. The ITS2 region was first amplified from molecularly confirmed species S using high fidelity Taq polymerase. The PCR reaction mixture (20 µL) contained 0.4 µM of primers 5.8S-F and ITS2C, 0.50 unit of Phusion® High-Fidelity DNA Polymerase, Phusion® HF reaction buffer (with 200 µM each dNTP and 1.5 mM MgCl_2_) and 0.5 µL of gDNA. The PCR thermal cycling protocol was: initial denaturation at 98˚C for 30 sec, 35 cycles of each of denaturation at 98 °C for 10 sec, annealing at 65 °C for 30 sec and extension at 72 °C for 30 sec, followed by a cycle of final extension at 72 °C for 7 min. The PCR product was purified using QIAquick PCR purification kit (Qiagen Inc, USA). Approximately 50 ng of PCR product was incubated at 70 °C for 20 min in a reaction mixture containing buffer, 1.5 mM of MgCl_2_, 0.5 unit of Taq DNA polymerase and 200 µM of dATP to incorporate ‘A’-tail at 3' end of the PCR product. The product was cloned in pGemT easy vector following Mishra *et al.* (2021). A total of 10 clones were sequenced from both directions of the strand.

**Specificity of T-specific primer**

Two PCR assays were performed to check the specificity of species T-specific primer. In the first PCR 10 of each of the *An. fluviatilis* S and T, 10 samples of *An. minimus,* as identified by DNA sequencing, and five ITS2-clonal samples of species S were subjected to ITS2 based species-diagnostic PCR (ITS2-ASPCR) developed by Manonmani *et al.* (2001). For convenience, we named the primers used in ITS2-ASPCR as 5.8S-F (universal forward primer), T-R (species T-specific, reverse) and S-R (species S-specific, reverse). Clones with correct and PCR-error-free insert (confirmed through DNA sequencing) were used as template DNA. In the second PCR, the ITS2-clonal samples of species S were amplified with primers 5.8S-F and T-R to check the specificity of T-specific primer (T-R). Two of the amplified products of the second PCR were sequenced from both directions of the strands to confirm that the mispriming of the primer T-R with species S is not due to contamination of reagents, carryover of DNA, or aerosol DNA. In both PCR assays, similar PCR conditions were used as described above. The stringency of PCRs was ensured by opting for minimum recommended MgCl_2_ concentration (1.5 mM), use of Hotstart Taq polymerase (0.5 unit), and keeping high annealing temperature (55 °C) to ensure that mispriming is not due to liberal PCR conditions.

**PCR-RFLP**

While species-specific PCR primers may be prone to non-specific extension, PCR-RFLP manifests tremendous sequence-specificity to their recognition sequence. We, therefore, designed a PCR-RFLP assay for the identification of species T based on ITS2 sequences to verify the presence of true hybridization in samples that showed prominent species S- and faint T- specific bands in ITS2-ASPCR. The unique species-specific restriction enzymes for species T were identified using an online tool available at <http://insilico.ehu.es/restriction/two_seq>. A unique restriction enzyme BtsI-v2 (GCAGTG_NN') specific to species T was used for PCR-RFLP. The amplified ITS2 products of 10 samples showing the presence of species S and T bands in ITS2-ASPCR were subjected to PCR-RFLP. For each PCR-RFLP, five μL of ITS2 product (amplified with primers ITS2A and ITS2C) was subjected to restriction digestion with five units of restriction enzyme in a reaction mixture (20 μl) and incubated at 37 °C for 4 hours. The reaction was heat-inactivated at 65°C for 20 min. The restriction products were run on a 2% agarose gel containing ethidium bromide and visualized under UV illumination.
